# A method for sampling parasitized *Drosophila suzukii* puparia from soil

**DOI:** 10.1101/2023.10.23.563656

**Authors:** Clarissa Capko, Jason Thiessen, Lana Harach, Jessica L. Fraser, Michelle T. Franklin, Paul K. Abram

**Affiliations:** Department of Biology, University of the Fraser Valley, Abbotsford, British Columbia V2S 7M8, Canada; Agriculture and Agri-Food Canada, Agassiz Research and Development Centre, Agassiz V0M 1A2, British Columbia; Department of Agriculture, University of the Fraser Valley, Chilliwack, British Columbia V2R 5X6, Canada; Université Laval, Centre de recherche et d’innovation sur les végétaux (CRIV), Pavillon Envirotron, Québec, QC G1V 0A6 Canada

**Keywords:** Insect soil sampling, overwintering parasitoids, biological control, spotted wing drosophila

## Abstract

Methods to measure the diversity and biological control impact of parasitoids for the control of spotted-wing drosophila, *Drosophila suzukii* (Matsumura) (Diptera: Drosophilidae) are being developed in support of biological control programs around the world. Existing methods to determine parasitism levels and parasitoid species composition focus on sampling *D. suzukii* within fresh and rotting fruit. However, many *D. suzukii* pupate in the soil or in dropped fruit, where additional parasitism could occur and where their parasitoids are thought to overwinter. Here we introduce a method for extracting parasitized *D. suzukii* puparia from the soil through a sieve and flotation system, allowing for effective collection of puparia, from which parasitoids can then be reared. Although the method considerably underestimates the absolute number of puparia in soil samples, it nonetheless yields a high number of puparia relative to sampling effort and provides a robust estimate of the relative abundance of puparia among samples. Using this method, we confirmed that at least five species of parasitoids, including some that have rarely been detected in past studies, overwinter in their immature stages inside *D. suzukii* puparia in south coastal British Columbia, Canada. The ability to sample puparia from the soil will lead to a more comprehensive view of both *D. suzukii* and parasitoid abundance throughout the season, help confirm parasitoid establishment following intentional releases, and provide a way to measure the diversity of parasitoid species and potential interactions among parasitoids (e.g., hyper- or klepto-parasitism) that can often occur on the soil surface.

## Introduction

Spotted wing drosophila, *Drosophila suzukii* (Matsumura) (Diptera: Drosophilidae), is a vinegar fly native to southeast Asia that has invaded Europe, North and South America, and Africa (Asplen et al. 2015; Boughdad et al. 2021), causing major agricultural losses in several soft fruit crops (Tait et al. 2021). *Drosophila suzukii* lays its eggs in ripe fruits where the larvae develop and cause fruit to rot prematurely and become unmarketable.

Biological control methods have been the focus of much research since the invasion of *D. suzukii*, as parasitoids are better able to attack larvae within fruit compared to pesticides and could help to reduce *D. suzukii* populations across the landscape, including in unmanaged ecosystems that serve as sources for populations that infest soft fruit crops. Research initially focused on parasitoid species that were already present in the invaded areas, finding a few species that could help in managing *D. suzukii* with additional releases supporting the natural parasitoid populations (Wang et al. 2020). Further explorations of parasitoids of *D. suzukii* in its native range found that two larval parasitoids, *Ganaspis brasiliensis* (Ihering) (Hymenoptera: Figitidae) and *Leptopilina japonica* (Novković & Kimura) (Hymenoptera: Figitidae), were the most abundant and these species were subsequently tested for host specificity in quarantine laboratories (reviewed in Wang et al. 2020). *Ganaspis brasiliensis* has proven to be the most host specific species (reviewed in Wang et al. 2020) and releases have taken place in Italy (Fellin et al. 2023) and are ongoing in the United States (X. Wang, personal communication). Prior to these intentional releases however, one or both of *L. japonica* and *G. brasiliensis* were found to have established in some parts of Canada, the United States and Europe (Abram et al. 2020, Beers et al. 2022, Puppato et al. 2020).

The seasonal ecology of both *G. brasiliensis* and *L. japonica* during the growing season has previously been documented in British Columbia, Canada (Abram et al. 2022a) and involved collections of *D. suzukii* larvae and/or puparia in fruit found on host plants, from which parasitoids were then reared (Abram et al. 2022b). However, *D. suzukii* readily pupates in the soil and in fruit that has dropped to the soil surface (Woltz & Lee, 2017), and additional parasitism or hyperparasitism may take place on the soil (Wang et al. 2016, Haussling et al. 2022). In addition, overwintering of parasitoids of *D. suzukii* is thought to take place in their immature stages inside host puparia (Hougardy et al. 2019), which at the end of the growing season would be primarily located on or just under the soil surface. The overwintering biology of *D. suzukii* parasitoids has only been studied under laboratory conditions (Wang et al. 2018, Amiresmaeili et al. 2020, Häner et al. 2022), but field evidence that *D. suzukii* parasitoids do in fact overwinter inside host puparia is lacking.

To study parasitism of *D. suzukii* puparia in the soil, an efficient sampling method to collect puparia is needed. Methods have been developed to sample various dipteran puparia to measure both fly and parasitoid abundance and diversity in the reproductive and overwintering seasons (Harcourt & Binns, 1980, Maier, 1981, Turnock et al. 1995). These methods typically involve the use of a single sieve to remove puparia from soil samples (Maier, 1981, Turncock et al. 1995, Biancheri et al. 2022), with flotation in a beaker of water as an additional step (Harcourt & Binns, 1980). A new, tailored, and efficient method of sampling overwintering Drosophilidae parasitoids from the soil would help provide insight into parasitism levels and parasitoid species composition throughout the season, the timing of parasitoid diapause in the fall, and the timing of spring emergence in the field of *D. suzukii* parasitoids, including both adventive and native species. Also, with the releases of parasitoids for biocontrol and increasing detections of adventive populations in new areas, effective methods to monitor population and community dynamics will be useful to understand the biological impacts of parasitoids on *D. suzukii*. However, only one study, to our knowledge, has developed methods to sample parasitized Drosophilidae (including *D. suzukii*) puparia from the soil (Biancheri et al. 2022). None of these methods have been validated or presented in such a way that they can be replicated by other researchers.

In this study, we developed a method for sampling *D. suzukii* puparia from the soil. The mechanism involves a cascading multi-sieve system along with a floating technique to efficiently separate few or many puparia from soil samples, building on methods developed for various invertebrate and dipteran species (Harcourt & Binns, 1980, Doane et al. 1987, Edwards, 1991, Chavalle et al. 2015). We demonstrate that it is possible to use this method to determine: (1) whether the adventive larval parasitoids of *D. suzukii* (*L. japonica* and *G. brasiliensis*) overwinter in host puparia in the soil in immature stages; (2) the parasitoid species composition in overwintering *D. suzukii* puparia; and (3) the development time of these parasitoids in the laboratory and a corresponding prediction regarding their timing of emergence in the spring.

## System Design

The flotation system was made from three 20 L plastic buckets, 3 irrigation tubes, a 4.76 mm-mesh sieve, a 3.35 mm-mesh sieve, a 1 mm-mesh sieve, and a 250 µm-mesh sieve (Figure 1). The materials and cost to construct the flotation system are listed in Table 1.

**Figure 1.**
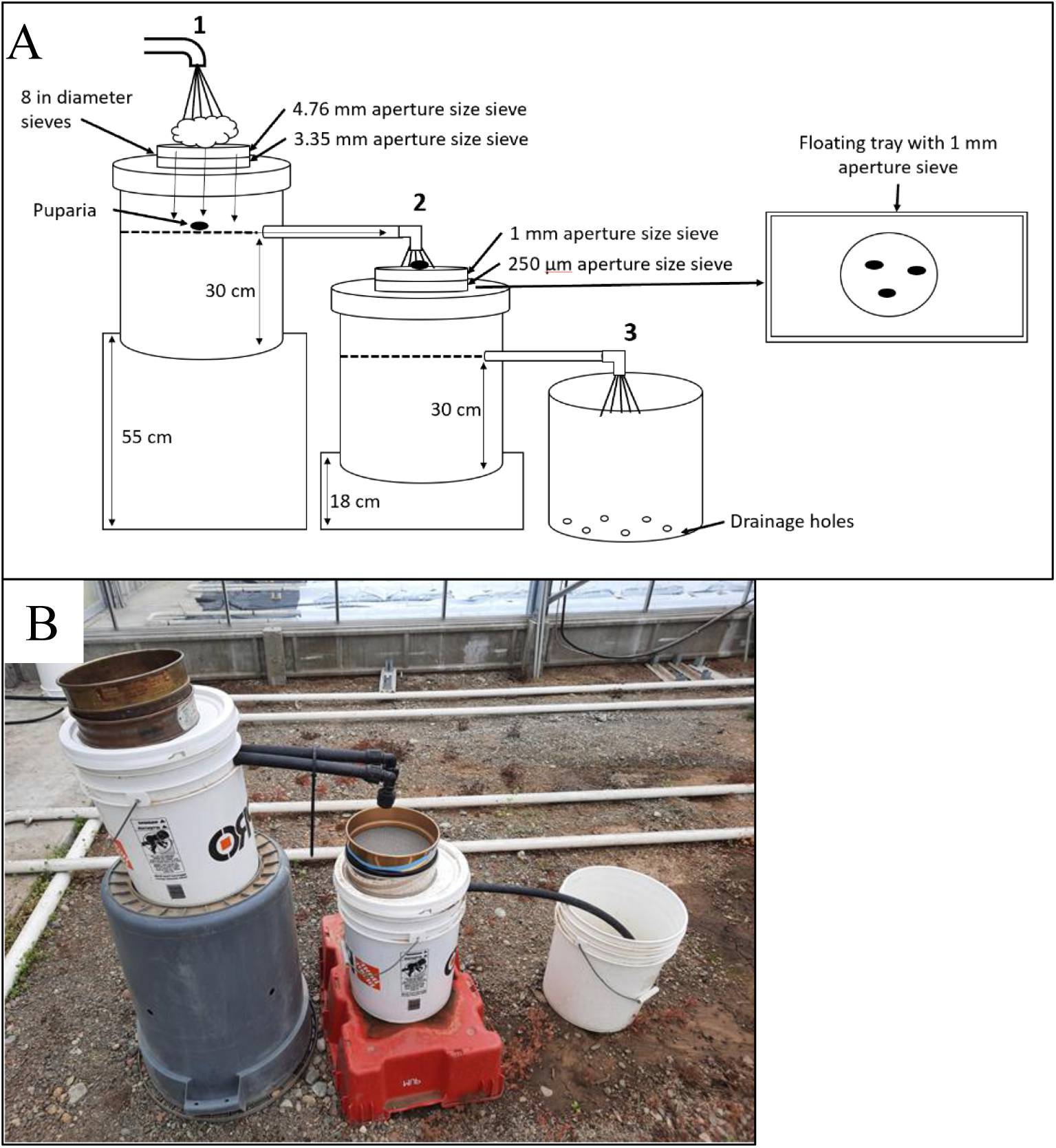
A. Bucket flotation system to separate out floating puparia. Two irrigation tubes were set into bucket 1 roughly 7 cm below the lip with elbows for water to flow downwards into bucket 2. One irrigation tube was set into bucket 2 at the same height. Holes in the bucket lids were made for the sieves to sit in. Bucket 1 had the 3.35 mm sieve set first on the lid, with the 4.76 mm sieve on top. Bucket 2 had the 250 um sieve set first on the lid, with the 1 mm sieve on top. Bucket 1 was on a 55 cm bin off the ground, bucket 2 was placed 18 cm off the ground, and bucket 3 was placed on the ground with holes drilled in the bottom to allow water to flow out. B. Image of bucket flotation system used in this study.

**Table 1.** Description of the materials, suppliers, and estimated costs for the construction of the flotation system. Total cost of the set up is estimated at $547.35 CAD, excluding filter paper and petri dishes. Prices quoted are per individual item.

| Supply | Description | Supplier | Estimated Cost | Product link |
| --- | --- | --- | --- | --- |
| Three Plastic Buckets | Plastic Food Grade Safe Bucket, 19 L, White | Canadian Tire | \$4.99 | https://www.canadiantire.ca/en/pdp/canadian-tire-plastic-food-grade-safe-bucket-5-gal-19-l-0581060p.0581060.html |
| Three Bucket Lids | Plastic Food Grade Safe Bucket Lid, 19 L, White | Canadian Tire | \$3.29 | https://www.canadiantire.ca/en/pdp/canadian-tire-plastic-food-grade-safe-bucket-lid-5-gallon-0581031p.0581031.html?rrecName=Similar%20Items%20&rrecReferrer=product&rrecProductId=0581031P&rrecProductSlot=1&rrecSchemeId=product1_rr&rrec=true |
| No.4 4.76 mm sieve | Stainless steel 8 in full sieve height, standard grade | W.S. Tyler | \$91.69 | https://advantechmfg.com/stainless-steel-stainless-steel-8-astm-test-sieve/ |
| No.6 3.35 mm sieve | Stainless steel 8 in full sieve height, standard grade | W.S. Tyler | \$91.69 | https://advantechmfg.com/stainless-steel-stainless-steel-8-astm-test-sieve/ |
| No.18 1 mm sieve | Stainless steel 8 in full sieve height, standard grade | W.S. Tyler | \$91.69 | https://advantechmfg.com/stainless-steel-stainless-steel-8-astm-test-sieve/ |
| No.60 250 µm sieve | Stainless steel 8 in full sieve height, standard grade | W.S. Tyler | \$91.69 | https://advantechmfg.com/stainless-steel-stainless-steel-8-astm-test-sieve/ |
| Hand valve | Brass one touch valve | Dramm | \$29.85 | https://canadagrowsupplies.com/products/dramm-brass-one-touch-valve |
| Water Breaker | 1000PL nozzle | Dramm | \$24.85 | https://canadagrowsupplies.com/products/dramm-1000pl-redhead-water-breaker-nozzle |
| Irrigation tubes | ¾" x 100' 75PSI Standard Poly Pipe | Home Hardware | \$54.99 | https://www.homehardware.ca/en/34-x-100-poly-pipe-75psi-standard/p/3250449 |
| White Tray | Laboratory tray, PP white, 300 x 400 mm, 10 L | Thomas Scientific | \$39.47 | https://www.thomassci.com/Laboratory-Supplies/Trays/_/Laboratory-Trays |
| Fine paintbrush | Synthetic gold sable round brush set | The Gilded Rabbit | \$6.59 | https://www.thegildedrabbit.ca/heinz-jordan-brush-series-700-gold-sable-round.html |
| Petri dish | Clear 50 mm diameter dish | Fisher Scientific | \$1.44 | https://www.fishersci.ca/shop/products/falcon-tight-fit-lid-dishes/08757105#?keyword=falcon%20dishes |
| Filter Paper | 45 mm standard grade, circle | VWR | \$0.08 | https://ca.vwr.com/store/product?keyword=C A76423-768 |

## Procedure

### Field sample collections

To obtain *D. suzukii* puparia that could contain overwintering parasitoids, 60 soil samples were collected from a cultivated blackberry (*Rubus fruticosis* L.) plot at the Agassiz Research and Development Centre in Agassiz, British Columbia (GPS: 49.24°, -121.75°). Twenty samples were collected on each of three dates in 2021: March 16, 23, and 30. These dates are early enough in the season that the parasitoids were not expected to be developing based on a minimum developmental threshold of 8.2 °C for *G. brasiliensis* and 10.5°C for *L. japonica* (Hougardy et al. 2019) and the average outdoor temperatures remaining below this threshold before March 16^th^ (Environment and Climate Change Canada, 2023). After collection, samples were placed in a 5°C refrigerator (for up to 2 months) until processing. Any viable insects inside puparia were expected to be parasitoids, and not developing *D. suzukii*, as *D. suzukii* overwinters as adults (Zerulla et al. 2015, Wallingford et al. 2016).

To obtain soil samples with similar exposed surface area and total weight, we removed the screen from a cylindrical sieve (height = 6.7 cm diameter = 21 cm) and marked a 3 cm depth around its inside. To take each individual sample, the hollowed-out sieve was pushed into the soil to the 3 cm depth marker and soil was removed, using a hand trowel, and placed in a sealable plastic bag. The sample depth of 3 cm was considered to be adequate because *D. suzukii* do not typically pupate below 4 mm (Kanzawa, 1939). Because obtaining representative measurements of the absolute number of puparia per unit soil surface area or weight was not a goal of this study, the selection of sampling locations was non-random; they were taken from locations within the field where parasitoids were likely to be found based on soil texture (silt or loam rather than heavy clay) and fruit seed abundance.

### Floating procedure for field-collected samples

Before starting the process, the first bucket was filled with water, up to the level of the drain tubes, to help move the puparia to the second set of sieves. The weight of the soil sample was measured with a balance and then the soil was deposited onto the 4.76 mm-mesh sieve. A hose with running water, fitted with a hand valve for regulating flow and a water breaker to fan out the water was used to rinse the soil through. Any larger clumps of soil were broken up by hand, until only large non-soil particles were left behind which were rinsed and then discarded. The first sieve was then removed, and the water used again to rinse the soil through the 3.35 mm-mesh sieve. The lid of the bucket was then removed and any particles and puparia on the lid and inner walls of the bucket were rinsed into the water. Using the hose, puparia were rinsed through the irrigation tubes to the next set of sieves. The 1 mm and 250 µm-mesh sieves were then removed and individually placed in a white plastic tray (45 × 30 × 6cm) with 3-4 cm (depth) of water. Floating puparia were then removed using a fine paintbrush and placed into a Petri dish (diameter = 6 cm) lined with filter paper. Any broken or clearly empty puparia were removed. When the status of a puparium was unclear, it was observed under a dissecting microscope to search for the light-colored mass inside the puparium which indicates the presence of an immature parasitoid (C. Capko, personal observations). The time to process each sample, beginning with the weighing of the soil sample and ending when all puparia from a sample were in a Petri dish, was recorded.

### Parasitoid emergence from field-collected samples

Each puparium was individually identified as either a *D. suzukii* puparia or that of an unknown (other Drosophilidae) species using spiracular morphology (Abram et al. 2022b). Puparia that could not be assigned to species or that had broken spiracles (n = 57) were not included in the study’s analysis of parasitoid species composition. *D. suzukii* puparia were placed in Petri dishes with a maximum of 30 puparia per dish or individually in Eppendorf tubes. These were then placed in an incubator at 21.25°C and checked 3 times weekly for emergence (n= 1,302). An additional replicate of soil samples was done to set aside puparia (n = 490) for molecular analysis and were not included in this study. Parasitoids that emerged from the Eppendorf tubes and Petri dishes (n=727) were initially identified following Abram et al. (2022b) and representative vouchers were inspected to confirm identification by Dr. Matt Buffington (Figitidae), Dr. Robert Kula (Braconidae), and Dr. Michael Gates (Pteromalidae) at the Systematic Entomology Laboratory, USDA-ARS, National Museum of Natural History, Smithsonian Institution, Washington, D.C., USA. Voucher specimens are stored at the Agassiz Research and Development Centre, Agassiz, BC. Development time of each puparium from the Petri dishes only for species *L. japonica*, *G. brasiliensis*, and *Asobara cf. rufescens* (Förster) (Hymenoptera: Braconidae) (n= 550) was set as the midpoint between the date of emergence and the previous observation. Predicted emergence dates for *L. japonica* and *G. brasiliensis* were calculated using weather data extracted from the Environmental and Climate Change Canada Historical Climate Data website (https://climate.weather.gc.ca/index_e.html) from Agassiz Research and Development Centre Station, Agassiz, British Columbia for the years 2017-2022, along with estimates of lower thermal thresholds for parasitoid development from Hougardy et al. (2019) (See Results).

### Validation of floating technique

To test the recovery rate of puparia from soil samples, and to what degree the relative number of puparia extracted from a sample correlated with the true number of puparia it contained, we ran a validation experiment. Thirty samples of soil (approx. 950 g each) from a plot located within

500m of our initial soil sample collection sites, but where *D. suzukii* puparia were not expected to be present (because there were no fruit hosts grown there), were collected into plastic bags. Five of the 30 samples were left unaltered as “blank” negative controls. The remaining 25 samples were seeded with between 1 and 124 *D. suzukii* puparia containing *L. japonica* pre-pupae (total = 665; mean = 26.6). The numbers were randomly drawn from actual numbers extracted from real soil samples described above. Puparia containing parasitoid pre-pupae were reared by: (i) placing 20-30 mixed-sex adult *D. suzukii* (14-28 days since emergence) in vials with approximately 20 mL of potato flake diet (Formula 4-24 Instant Drosophila Medium, Carolina Biological Supply Company, USA) and half of a store-bought blueberry for 48h; (ii) removing flies and exposing immature *D. suzukii* to three mated female *L. japonica* for 2-3 days; (iii) removing parasitoids and incubating vials at 23°C, 16:8 h photoperiod, 50-70% RH for 14 days (at which point parasitoids were in the pre-pupal stage, the same stage at which they overwinter). Vials were stored at 8°C to arrest parasitoid development; parasitized puparia were then extracted from vials and counted by floating the samples in water and visually inspecting puparia for parasitism. A known number of parasitized puparia (see above) was then added to each of the soil samples, which were mixed and then stored at 5°C until they were processed using the flotation apparatus in the same way as the field-collected samples 10-14 days later. The investigator floating the samples was unaware as to the true number of puparia each sample contained. The number of puparia extracted from each sample was correlated with the known number that was originally added to the soil samples.

### Statistical analyses

Pearson’s correlation analyses and the production of graphics were done in R version 4.2.3 (R Core Team, 2023).

## Results

### Field-collected puparia

A total of 1,302 *D. suzukii* puparia were collected from soil samples to be reared (Table 2), with an average of 29.4 ± 4.4 (mean ± SE) (minimum = 0, maximum = 147) puparia extracted from each sample. The time required to process samples ranged from 8 to 38 min and was positively correlated with the number of puparia that were extracted from each sample (Pearson’s correlation; t = 11.2, df = 58, *r =* 0.83, p < 0.0001) (Figure 2).

**Figure 2.**
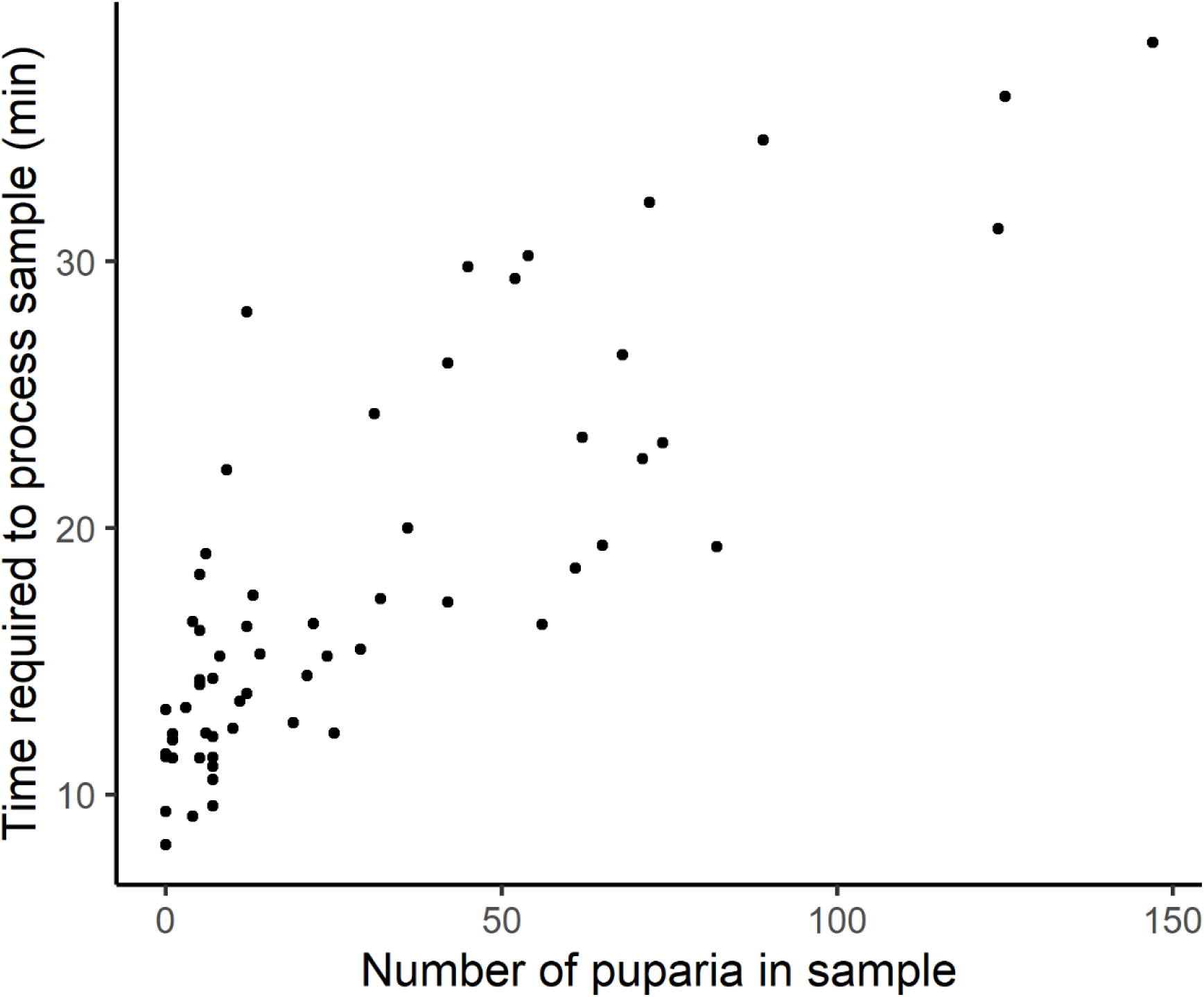
The time required to process each field-collected soil sample in relation to the number of puparia retrieved from each sample.

727 individuals of six different species of *D. suzukii* parasitoids emerged from the *D. suzukii* puparia extracted from soil samples (Table 2); the rest of the puparia did not emerge or have discernible contents. The most common parasitoid was *L. japonica* (90%) followed by *A. cf. rufescens* (6.5%) and *G. brasiliensis* (2.3%). The other parasitoids that emerged from puparia included *Spalangia erythromera* (Förster) (Hymenoptera: Pteromalidae), *Pachycrepoideus vindemiae* (Rondani) (Hymenoptera: Pteromalidae), and an unknown species belonging to the subfamily Pteromalinae (Hymenoptera: Pteromalidae) (M. Gates, personal communication).

**Table 2.** Summary of the number and species composition of parasitoids reared from *D. suzukii* puparia collected from the soil, the mean number of parasitoids recovered per sample, and the number of puparia from which no parasitoids emerged.

| **Species** | **Total number of puparia with each parasitoid species** | **Proportion of all emerging parasitoids** | **Mean number of parasitoids per sample** |
| --- | --- | --- | --- |
| *Leptopilina japonica* | 649 | 0.89 | 16.19 |
| *Asobara c.f. rufescens* | 44 | 0.065 | 1.61 |
| *Ganaspis brasiliensis* | 16 | 0.022 | 0.32 |
| Unidentified Pteromalidae | 10 | 0.014 | 0.322 |
| *Spalangia erythromera* | 6 | 0.0081 | 0.065 |
| *Pachycrepoideus vindemiae* | 2 | 0.00027 | 0.097 |
| Unemerged puparia | 575 | --- | --- |

When the puparia extracted from soil samples were reared in an incubator at 21.25 °C, *G. brasiliensis* adults took approximately 5 days longer to emerge than *A. cf. rufescens* and *L. japonica*, which had similar median development times (Figure 3). Degree day estimates for *L. japonica* were 232.1 (minimum =188.1, maximum = 553.0) and estimates for *G. brasiliensis* were 336.7 (minimum = 293.6, maximum = 400.0) (Table 3). Based on these ranges of development times and subsequent temperature data for the field for the past six years, predicted emergence dates would range from May 26 to June 29 (median) for *L. japonica* and *G. brasiliensis* (Table 3).

**Figure 3.**
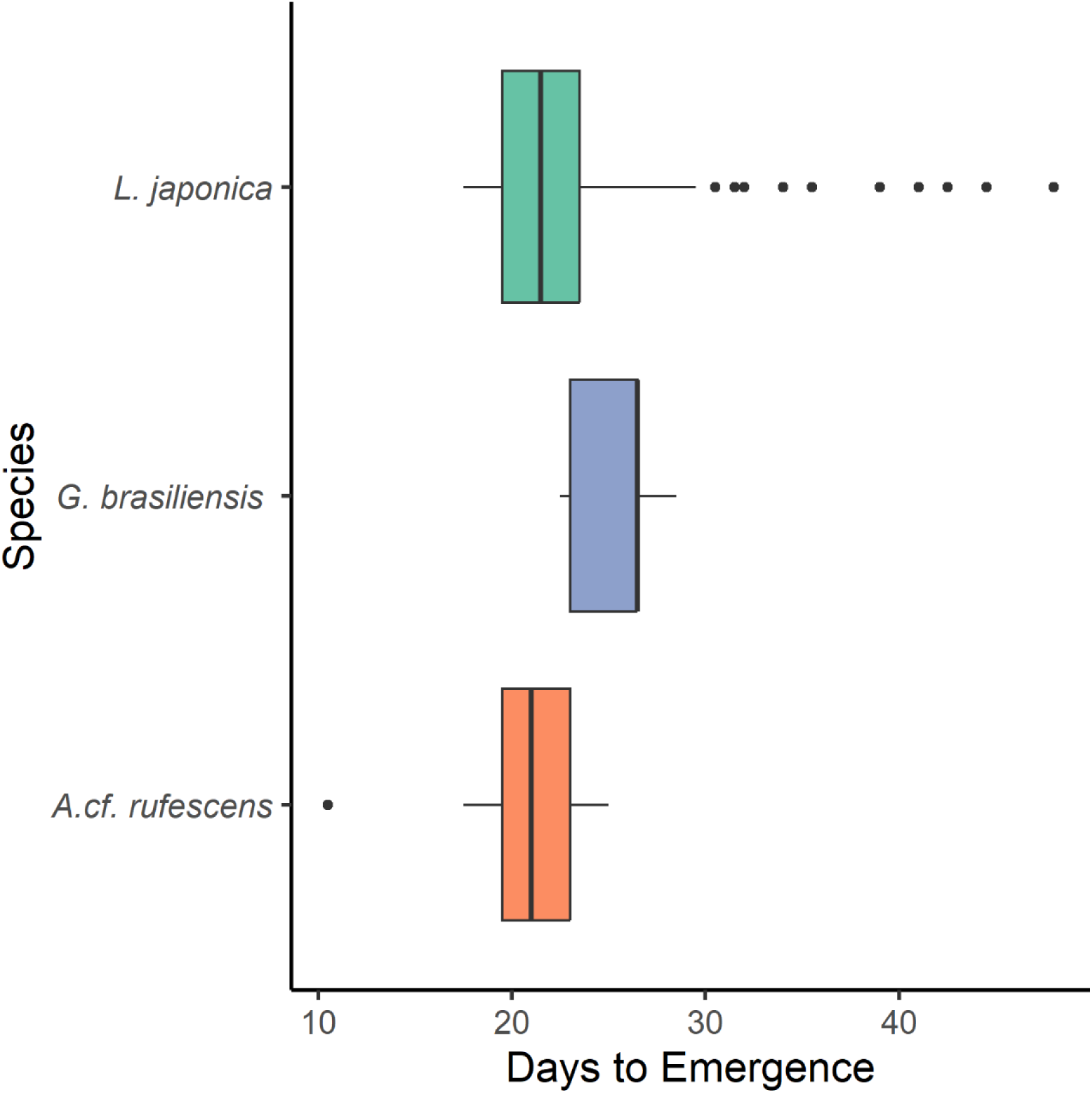
Development time of overwintering *Leptopilina japonica* (n = 501), *Ganaspis brasiliensis* (n = 13), and *Asobara* cf. *rufescens* (n = 36) reared from soil-collected puparia collected from the field and then kept at 21.25 °C. Lower and upper hinges indicate the first and third quartiles and any data greater or less than 1.5 times the inter-quartile range are shown as outlying data points. The median is displayed as the midline.

**Table 3.** Range of predicted emergence dates for *Leptopilina japonica* and *Ganaspis brasiliensis* using temperature data from years 2017-2022 from Agassiz, British Columbia, Canada. Estimates are based on the number of degree days required for overwintering immature parasitoids to complete their development and emerge in the lab at 21.25 °C using a base developmental temperature of 10.5°C for *L. japonica* and 8.2°C for *G. brasiliensis* (Hougardy et al, 2019).

| Species | Number of degree days required for diapausing parasitoids to emerge |  | Predicted Emergence Dates |  |  |  |  |  |
| --- | --- | --- | --- | --- | --- | --- | --- | --- |
|  |  |  | 2017 | 2018 | 2019 | 2020 | 2021 | 2022 |
| *Leptopilina japonica* | Minimum | 188.1 | June 8 | May 23 | May 26 | June 10 | June 3 | June 25 |
|  | Median | 232.1 | June 18 | June 2 | May 31 | June 19 | June 14 | June 29 |
|  | Maximum | 553.0 | July 21 | July 14 | July 10 | July 26 | July 12 | July 30 |
| *Ganaspis brasiliensis* | Minimum | 293.6 | June 6 | May 25 | May 21 | May 31 | June 2 | June 19 |
|  | Median | 336.7 | June 11 | May 30 | May 26 | June 9 | June 8 | June 24 |
|  | Maximum | 400.0 | June 20 | June 10 | June 2 | June 18 | June 17 | June 30 |

### Validation of floating method

The number of parasitized puparia recovered from samples by a blinded investigator was highly correlated with the number of parasitized puparia that had been added previously (Pearson’s correlation; r = 0.95, t = 16.4, df = 28, p < 0.0001) (Figure 4). There were no false positives; that is, no puparia were recovered from any of the five “blank” soil samples. There was one false negative; a sample in which one puparium was placed but none were recovered. On average, 46.4 ± 5.0% (mean ± SE) of puparia were recovered from each sample.

**Figure 4.**
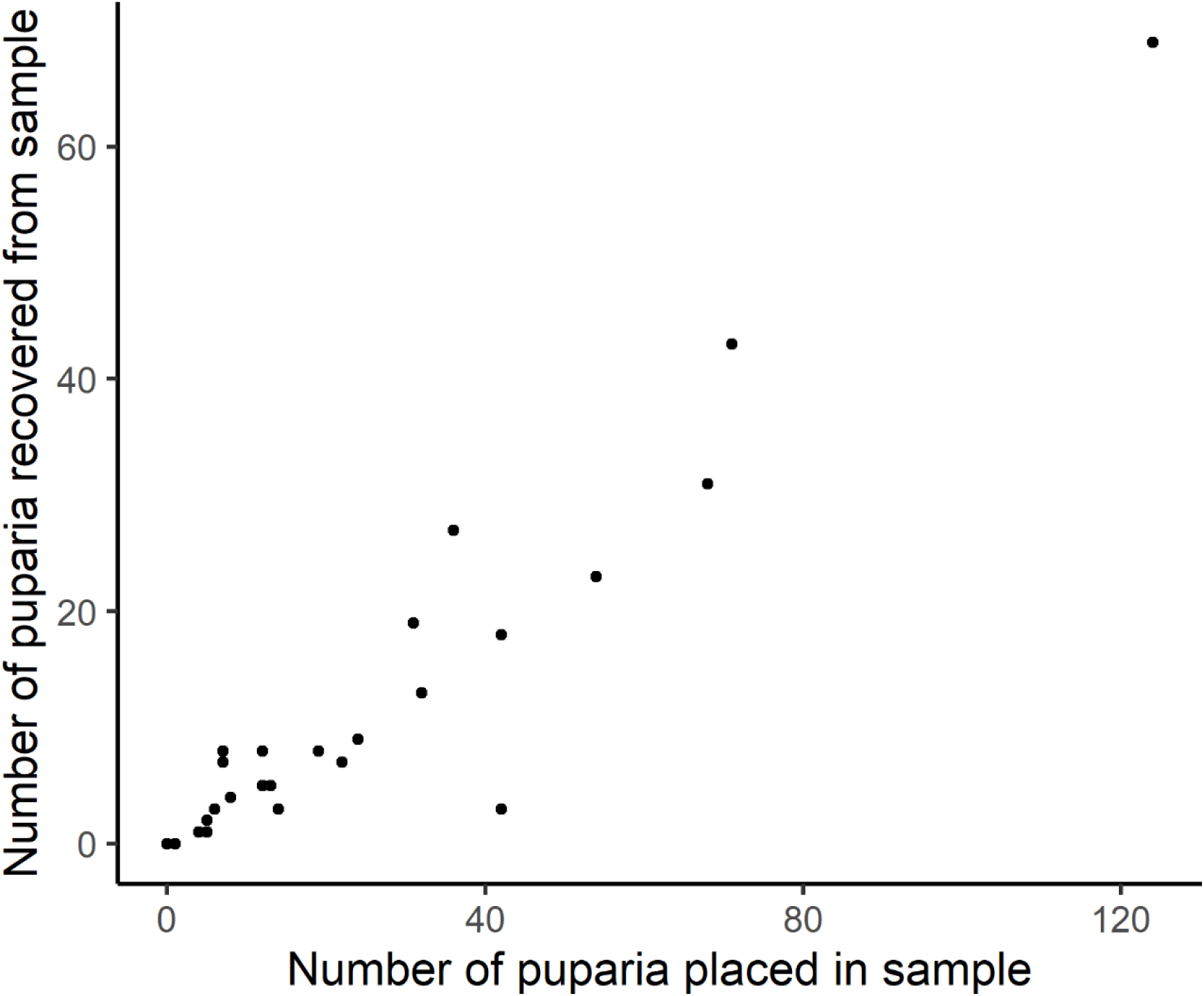
The number of parasitized puparia recovered from soil samples by a blinded observer plotted against the number of parasitized puparia that were placed in the sample by another investigator.

## Discussion

Our method was successful at removing parasitized *D. suzukii* puparia from the soil to determine parasitoid species compositions and relative abundance. The validation experiment indicated that while the method considerably underestimates the absolute number of puparia in soil samples, it does provide a very good indication of the relative number of puparia among samples. The latter result indicates that it will be useful for many research applications; indeed, most insect sampling techniques measure relative rather than absolute abundances. Using our soil sampling method to collect parasitized *D. suzukii* puparia during the winter in British Columbia, Canada, we confirmed that spotted wing drosophila parasitoids, both native and adventive species, overwinter in their hosts puparia in immature stages. Five species were found from overwintering puparia at varying proportions, with *L. japonica* being the most abundant (86-89%) and lower proportions of *A. cf. rufescens, G. brasiliensis, P. vindemiae,* and *S. erythromera. Asobara. cf. rufescens*, *P. vindemiae*, and *S. erythromera* are typically rare in fruit sampling (Abram et al. 2020; 2022a), and are candidate kleptoparasitoids and hyperparasitoids (Kraaijeveld, 1999, Wang & Messing, 2004, Gibson, 2009). This finding may indicate that soil sampling has the potential to reveal different final outcomes of parasitoid community interactions than fruit sampling, because it can capture the result of additional parasitism events that occur on the soil surface.

Various methods to sample soil invertebrates have been used in the past, including brine solutions, reduced air pressure, and organic solvents which can be costly and difficult to use (Edwards, 1991). The only study to apply a method for removing *D. suzukii* puparia from the soil used a single 1mm mesh sieve (Biancheri et al. 2022), which we also used; however, we found it useful to add additional larger sieves to remove larger debris first while keeping the entire process in a single set-up. We also included the step of floating the puparia in water to make it easier to remove them from the sieve. For *Drosophila* puparia specifically, we found that floating in water was sufficient to remove puparia from the soil after informally comparing it with a brine solution, which did not improve recovery. One disadvantage to this set-up is that it does require a larger designated space (approximately 1 × 2 m) with adequate drainage, unlike smaller systems such as the one developed by Edwards and Heath (1962).

Although we did not set out to measure the per-unit-area relative abundance of parasitized *D. suzukii* puparia in the soil in our pilot study, our method could be used to accomplish this goal in future studies. If the density of parasitized puparia in the soil is high, the method can yield sufficient numbers of parasitoids to study species composition and factors influencing diapause induction. In our pilot study, we took advantage of a blackberry field known to be highly infested by *D. suzukii* with a high fruit density dropping onto an exposed soil surface, likely contributing to the considerable number and diversity of parasitoids we found. In many habitats where *D. suzukii* infests fruiting plants, low numbers of parasitized puparia and minimal exposed soil surface may introduce challenges in determining parasitoid species composition, diapause timing, or measures of parasitoid density without taking larger numbers of samples. To measure parasitoid species composition or diapause timing in settings with lower densities of naturally occurring puparia, we suggest placing a relatively high number of infested fruits on the ground in a designated location, and then returning to that exact location later to sample the soil to provide a larger sample size. This ‘fruit placement method’ would not be appropriate for attempting to measure relative parasitoid density per unit soil surface.

One advantage of being able to sample parasitoids during their overwintering stages is that they can then be reared to estimate the timing of their spring emergence. Under laboratory conditions at constant high temperatures, *L. japonica* took less time to develop to adulthood from its diapausing stage than *G. brasiliensis*, which is in line with previous laboratory studies of egg-adult development of the two species (Wang et al. 2018, Hougardy et al. 2019). However, because *G. brasiliensis* has a lower minimum temperature threshold for development, the predicted emergence timing of the two species in the field is predicted to be similar; this is supported by a past study showing that the two parasitoid species are first found parasitizing *D. suzukii* within a week of each other in British Columbia (Abram et al. 2022a). The predicted timing of parasitoid emergence in 2020 (Table 3), the year of the Abram et al. (2022a) study, also corresponds quite well to when the two species were first observed in that study which was conducted in the same region as the current work. However, these relative timings of spring emergence of the parasitoids will need to be validated by future studies.

This soil sampling method provides a way to assess the incidence, location, and relative (but not absolute) abundance of overwintering parasitoids of *D. suzukii* in the field and provides an additional method of monitoring parasitoid-host ecological relationships during the fly and parasitoids’ reproductive season. With current releases of parasitoids across the United States and Europe (Fellin et al. 2023, Seehausen et al. 2022), methods that allow for the assessment of parasitoid establishment can help to guide the number of parasitoids that need to be released each year and understand whether hyperparasitism by native larval and pupal parasitoid species (Hougardy et al. 2022) may be occurring on the soil surface that could affect the establishment or efficacy of released larval parasitoids. Future research into the parasitoids overwintering biology could help to better assess biological control success in different climates and guide decisions on which species or strains will be more effective. Sampling puparia from the soil and fruit during the growing season could provide new knowledge on which parasitoid species are better able to parasitize puparia on the soil surface, compared to puparia in fruit.

## Acknowledgements

We thank Markus Clodius and Seth Nussbaum for offering the use of the blackberry field sampled in this study. We thank Warren Wong for help with rearing. Funding to PKA was from Agriculture and Agri-Food Canada and the Organic Science Cluster, part of the Canadian Agriculture Partnership.

